# Lactate promotes cardiomyocyte dedifferentiation through metabolic reprogramming

**DOI:** 10.1101/2020.07.21.213736

**Authors:** Jesús Ordoño, Soledad Pérez-Amodio, Kristen Ball, Aitor Aguirre, Elisabeth Engel

## Abstract

Cardiomyocytes undergo different metabolic changes during development and differentiation crucial for their maturation and adult function, such as contraction, growth and survival. Alterations of cardiac metabolism have been associated with multiple disease states and pathological hypertrophy. The shift in substrate preference can impair the stress response, but it may also have a role in cell growth and survival. Here, we evaluated the response of cardiomyocytes to the presence of exogenous lactate, an important metabolite for the fetal heart and cardiogenesis. Lactate-exposed mouse primary cardiomyocytes and human iPSC-derived cardiomyocytes quickly acquired a characteristic dedifferentiated phenotype, with enhanced proliferative capacity as determined by an increased expression of cell cycle (Ki67) and cytokinesis (Aurora-B) effectors. This effect was specific to cardiomyocytes and did not affect other heart cell populations (e.g. cardiac fibroblasts). Nevertheless, cardiac fibroblasts exposed to lactate promoted a pro-regenerative environment through the modulation of the release of cytokines (such as Fas, IL-13 or SDF1a). We characterized lactate-induced cardiomyocyte dedifferentiation through RNA-sequencing and gene expression analysis and identified increased expression of BMP10 (a TGFβ family protein involved in embryonic cardiomyocyte proliferation and stemness) and proteins associated to cell fate regulation (LIN28, TCIM) together with downregulation of cardiac maturation genes (GRIK1, DGKK). Bottom-up analysis suggested the phenotype promoted by lactate could be related to the activation of hypoxia signaling pathways. This finding indicated that, indeed, lactate may be a key player of hypoxic regenerative responses in the heart, as it usually accumulates as a result of glycolysis. In addition, *ex vivo* neonatal heart culture showed prolonged beating function and cardiac tissue integrity when culture media was supplemented with lactate. Thus, we conclude that lactate enhances cardiac proliferation by reprogramming cardiomyocytes towards a dedifferentiated stem cell-like state, supporting the notion that modulation of the metabolic microenvironment might be a powerful novel approach for promoting cardiac regeneration and tissue engineering applications.

## INTRODUCTION

Cardiovascular disease (CVD) is the leading cause of death in the developed world. Myocardial infarction and cardiac failure, two prevalent forms of CVD, are among the most important causes of mortality and morbidity worldwide [1]. During myocardial infarction, a significant portion of the myocardium dies and is replaced by fibrotic tissue. The subsequent loss of cardiac contractility gradually leads to heart failure and eventually death. Therapeutic strategies with varying degrees of success have been proposed to stop this progression [2–4]. However, in spite of these efforts, cardiac regeneration and full functional recovery have not been achieved yet [5–7]. An emerging trend for tissue regeneration strategies is the manipulation of the metabolic microenvironment to induce stem cell activation [8–10].

The mammalian heart has a very high energy demand as it must contract incessantly to guarantee oxygen supply to all the organs in the body. For this reason, heart metabolism is highly flexible and can vary widely depending on energy substrate availability. Interestingly, substrate preference changes very significant throughout the life cycle [11]. In early development, the fetus resides in a low oxygen environment and the placenta generates a lactate-rich environment, making it an important substrate for fetal metabolism [12]. The fetal heart is highly dependent on glycolysis at these stages and uses lactate as a primary metabolic substrate [13]. Coincidentally, fetal cardiomyocytes are very proliferative and present a poorly differentiated phenotype when compared to adult cardiomyocytes. After birth, cardiomyocytes switch from hyperplasia to hypertrophy, ceasing proliferation and increasing cell size in an oxygen-rich environment. This change prompts a new metabolic microenvironment in which glycolysis and lactate levels are greatly reduced, while fatty acid β-oxidation increases to become the predominant energy source for mature cardiomyocytes. In this oxygen-rich environment cardiomyocytes acquire their definitive mature differentiated state [14]. Remarkably, cardiomyocytes increase their lactate uptake after heart failure, likely to attenuate the negative effects of hypertrophy [15]. Some clinical trials have already proven the capacity of lactate to improve heart performance after acute heart failure [16]. Despite these interesting findings, there is little to no mechanistic knowledge into the cardioprotective effects of lactate. However, cardiac regeneration studies have shown that during heart regeneration, dedifferentiated cardiomyocytes appear *de novo* from adult cardiomyocytes and contribute to replenishment of heart muscle through sustained proliferation [4,17]. These dedifferentiated cardiomyocytes shift to a glycolytic metabolism and are thus more reliant on lactate than mature cardiomyocytes [4].

Lactate produced from glycolysis can be a metabolic substrate for oxidative phosphorylation, but also a biochemical signal through its effects on the lactate-binding receptor and hypoxia signaling pathways. It has shown to play important roles in angiogenesis, accelerating wound healing and potentiating neurogenesis [18–20]. Lactate can also induce changes in gene expression contributing to metabolic reprogramming and the development of a dedifferentiated phenotype on many cell types, such as myotubes or immune cells, upregulating thousands of genes associated with stemness [19,21]. Nevertheless, the regulatory mechanisms of lactate signaling are still uncertain. Some studies suggest a direct relationship between lactate and hypoxia response, as a mechanism to ensure maximal growth in prolonged hypoxia [19,22].

Given these previous evidences, we hypothesized that exogenous lactate might be an important signaling molecule on cardiomyocyte maturation. We report that lactate induces dedifferentiation and proliferation of both mouse and human cardiomyocytes, while it activates a reparative program on cardiac fibroblasts, and propose a novel mechanism of action for its effects on cardiac tissues. We also found that lactate improves heart function without affecting fibrosis. Our data provide new roles for lactate in cardiovascular biology and cardiac stem cells, and paves the way for new regenerative medicine approaches employing lactate for the induction of regenerative programs in the failing myocardium.

## RESULTS

### Effect of lactate on neonatal cardiomyocyte survival and proliferation

To evaluate the possible toxicity and optimize the suitable concentration of lactate on neonatal mouse cardiomyocytes, we used different concentrations of L-lactate, from 0 to 20mM, and cells were evaluated periodically. Higher concentrations were not considered because of excessive media acidification (data not shown). After 7 days of incubation, cell death was almost undetected in a live/death assay, and lactate showed no effect on cell viability at all different concentrations tested (Figures 1A and S1A). The evaluation of extracellular LDH confirmed the lack of cell death caused by the addition of 20mM of lactate to cardiac cells, as compared to the control (Figure 1B). Cell death was very low until 9 days in culture, but it increased rapidly after this point, both in control and lactate media. Therefore, we used a 20mM concentration of L-lactate for the following experiments on neonatal mouse cardiac cells.

**Figure 1.**
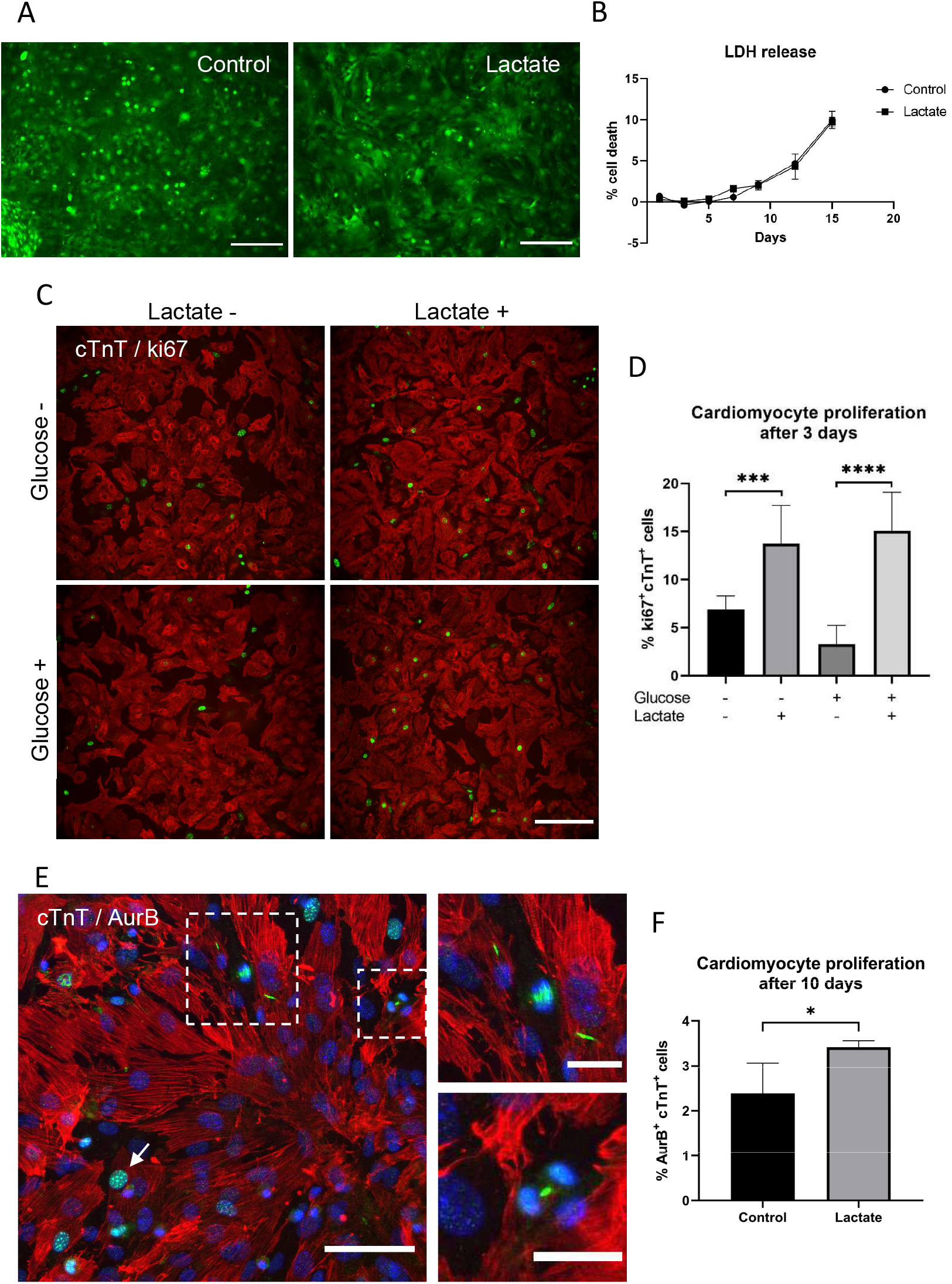
Lactate-induced cardiomyocyte proliferation. **(A)** Live/death assay of cardiomyocytes after 7 days of incubation with 0 (Control) or 20mM (Lactate) of supplemented L-lactate. Green: alive cells; red: dead cells; scale bar 200μm. **(B)** LDH detection on culture media of cardiomyocytes incubated with 0 (Control) or 20mM (Lactate) of supplemented L-lactate for 15 days. Data are expressed as % of cell death compared to maximum LDH released by lysed cells (100%). n = 3. **(C)** Ki67 immunostaining on cardiac cells after 3 days of culture with (+) or without (−) glucose or lactate. Red: cardiac Troponin T (cTnT), green: Ki67; scale bar: 150μm. **(D)** Quantification of ki67-positive cardiomyocytes after 3 days of culture with (+) or without (−) glucose and with or without lactate. Data are represented as % of ki67^+^cTnT^+^ within all cTnT^+^ cells. One-way ANOVA, ***p < 0.001, ****p < 0.0001; 10 random images and around 1200 cardiomyocytes were analyzed for each condition. **(E)** Aurora-B kinase immunostaining of cardiomyocytes after 6 days of culture. Dashed regions are magnified (right). Different mitotic stages can be identified, such as prophase (white arrow), anaphase and cytokinesis (top right magnification) or telophase (bottom right magnification). Green: AurB, red: cTnT, blue: nuclei; scale bar: 70μm (and 25μm for magnifications). **(F)** Quantification of AurB-positive cardiomyocytes after 10 days of culture without lactate (control) or with 20mM lactate-supplemented media. Data are represented as % of AurB+cTnT+ within all cTnT+ cells. Student’s T-test, *p < 0.05.; n = 4; at least 24 random images and 4000 cardiomyocytes were analyzed per for each condition.

Since the evaluation of cellular viability suggested a positive effect of lactate on mouse cardiomyocytes (Figure S1B), the proliferation of cardiomyocytes (cTnT^+^ cells) was evaluated. Lactate significantly increased the number of proliferating cells, as demonstrated by the increase in ki67^+^cTnT^+^ cells (Figure 1C and 1D). As glycolysis is the major energy source for proliferating cardiomyocytes [11], we tested whether glucose availability could influence the effect of lactate on cardiomyocyte proliferation. Figure 1D also shows that the increase in ki67^+^cTnT^+^ cells produced by lactate was not affected by the presence or absence of high concentrations of glucose (around 20mM), suggesting an independent pathway for the observed effect. A similar effect was observed when cTnT^-^ cells were not discriminated (Figure S1C). However, when lactate was removed from culture media after 3 days of incubation, the number of ki67^+^ cells decreased (Figure S1D), suggesting that lactate is continuously required for cell proliferation.

Multinucleation and polyploidy are characteristic traits of cardiomyocytes, and to further validate cardiomyocyte proliferation, the expression of Aurora-B kinase (ARK-2 or AurB), a key regulator for cytokinesis during mitosis, was also evaluated (Figure 1E). Consistently, the number of AurB^+^cTnT^+^ cells was significantly increased in the presence of lactate (Figure 1F).

### Effect of lactate on cardiac fibroblasts

Extensive cardiac fibrosis after a myocardial infarction is detrimental and may eventually lead to heart failure [23]. Therefore, we studied the non-myocytic cell fraction of isolated mouse hearts and evaluated the implications of lactate on fibrosis. Almost all cells from this fraction showed expression of vimentin, an intermediate filament highly expressed in fibroblasts (Figure 2A) and about half the population were also positive for alpha smooth muscle actin (α-sma), a characteristic phenotype of cardiac myofibroblasts (Figures 2B). The number of ki67^+^ cells was evaluated with lactate and/or glucose (Figure 2C). EdU (5-ethynyl-2’-deoxyuridine) incorporation was also assessed after incubation with different concentrations of lactate (Figure S2A and S2B). Contrary to cardiomyocytes, lactate showed no effect on the number of proliferating cells, regardless the presence of high concentrations of glucose. No significant differences were neither observed on collagen production after 3, 6 or 10 days of incubation with lactate compared to control (Figure 2D). A scratch wound assay was performed to evaluate migration of non-myocyte cardiac cells (Figure 2E). After 24 hours, cells incubated with and without lactate showed the same amount of recovered area, indicating a similar migratory behavior. The fraction of myofibroblasts present in the culture was assessed by flow cytometry (Figure S2C), indicating no significant difference in the number of vimentin and α-sma positive cells in the presence of lactate.

**Figure 2.**
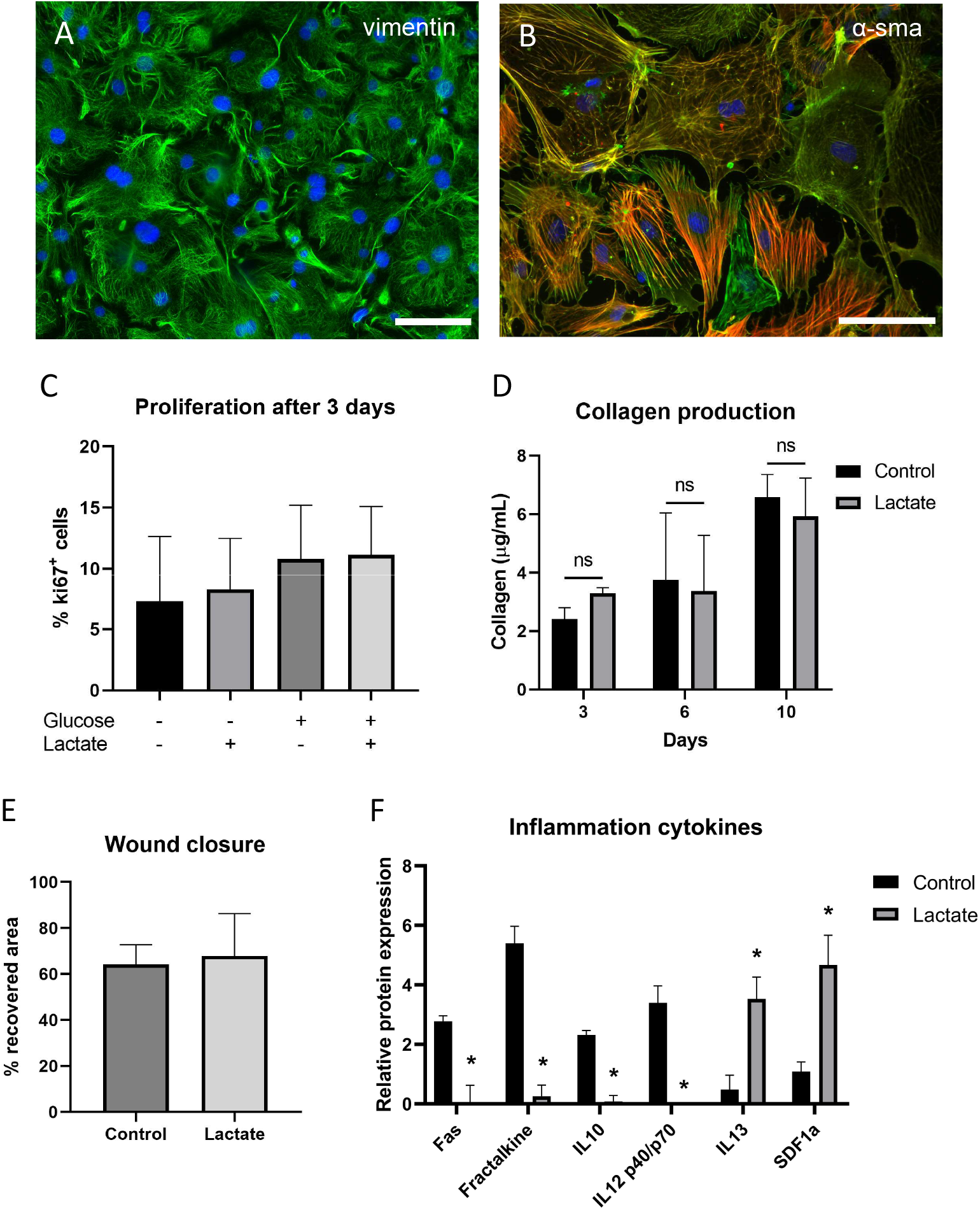
Effect of lactate on non-myocyte cells. **(A)** Vimentin immunostaining on non-myocyte cells. Green: vimentin, blue: DAPI; scale bar 150μm. **(B)** Alpha smooth muscle (α-sma) immunostaining on non-myocyte cells. Green: actin, red: α-sma, blue: DAPI; scale bar 100μm. **(C)** Ki67-positive cells quantification after 3 days of culture with (+) or without (−) glucose and with or without L-lactate. One-way ANOVA. Data are represented as % of ki67+ within all cells. 10 random images were analyzed for each condition. **(D)** Collagen quantification on cardiac fibroblasts after 3, 6 and 10 days of culture with or without 20mM of lactate. Two-way ANOVA, ns: non-significant. **(E)** Wound closure measured as % of recovered area after 24 hours of scratching with or without lactate. **(F)** Significantly upregulated and downregulated inflammatory cytokines from an inflammation antibody array of 40 cytokines. Signal expression is corrected by total protein amounts and normalized to internal positive (100) and negative controls (0). Student’s T-test of lactate compared to control condition, *p < 0.05.

In addition, the expression of different inflammatory cytokines was evaluated upon exposure to lactate (Table S1). Interestingly, lactate significantly reduced the expression of Fas, Fraktalkine, IL-10 and IL-12p40/p70, shown to be prejudicial for cardiac recovery after injury, while increased the expression of IL-13 and SDF1a, known to improve healing and remodeling after a myocardial infarction (Figure 2F).

Therefore, lactate did not show any effect on proliferation, collagen production, migration or myofibroblast transdifferentiation of the non-myocytic population of cardiac cells, mainly composed by cardiac fibroblasts, but appears to generate a pro-regenerative environment for cardiomyocytes, thus promoting cell survival and cardiac repair.

### Expression of progenitor and maturation genes

To further understand the molecular mechanisms by which lactate stimulates proliferation of mouse cardiomyocytes, we analyzed the expression of different progenitor genes by RT-qPCR (Figures 3A and S3A). We found that the continued supplementation with L-Lactate in cell media promoted an increasing expression of bone morphogenetic protein-10 (*Bmp10*) and transcription factor *P63*, reaching over a 4 times-fold expression after 10 days of culture compared to control without lactate (4.8 ± 0.95 and 4.2 ± 0.97 fold change respectively). Both genes have shown to be critical for cardiogenesis and the maintenance of a progenitor and proliferative phenotype of cardiomyocytes [24,25]. Additionally, the expression of *Tcim*, a positive regulator of the Wnt/β-catenin pathway associated to a poor differentiation state [26], was also increased in the presence of lactate (Figures 3A and S3A). In a lower extent, lactate also stimulated the expression of *Gata4* and *c-kit*, typically associated to cardiac progenitor cells.

**Figure 3.**
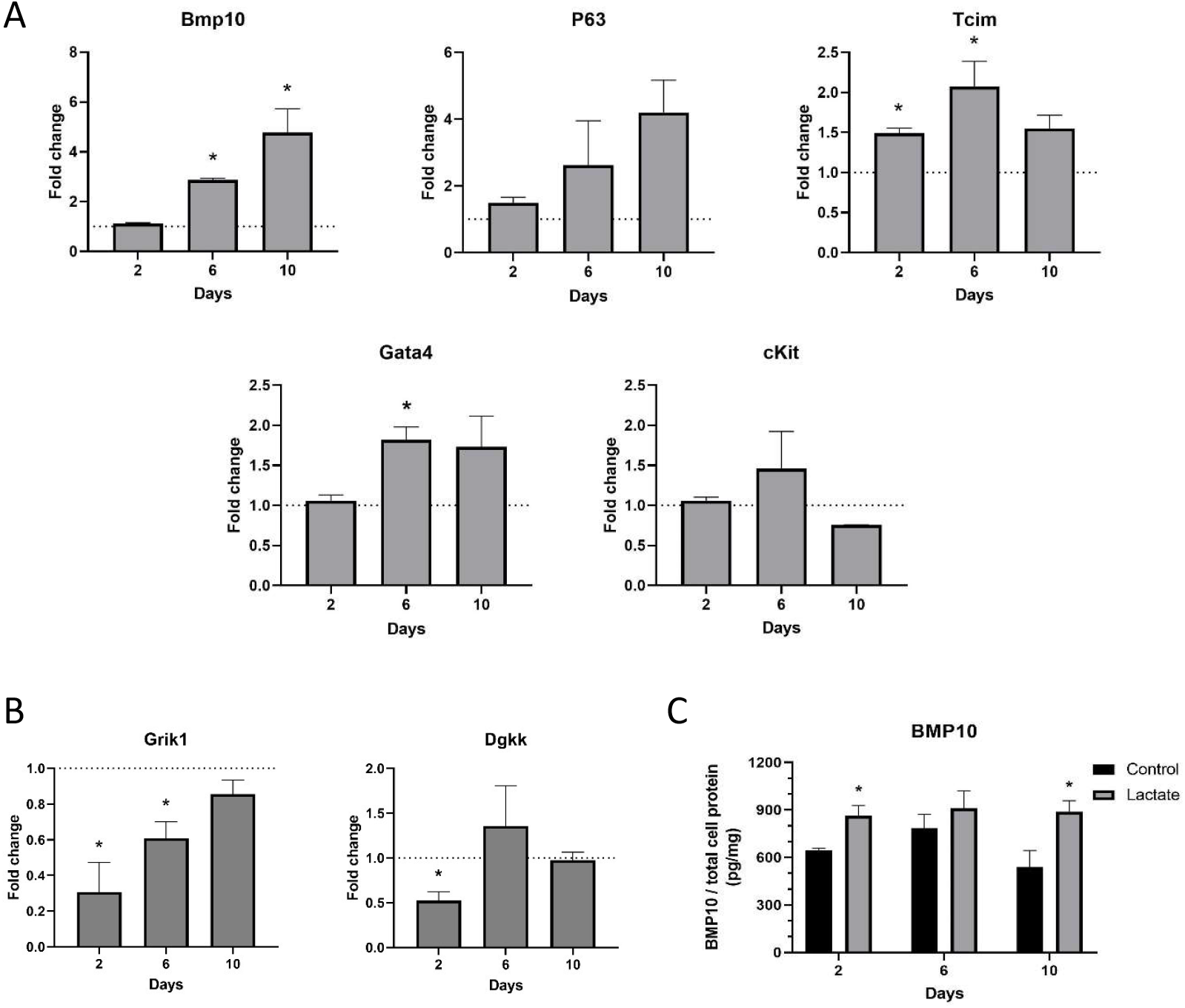
Gene expression of mouse cardiomyocytes. **(A)** Relative expression of progenitor genes in cardiomyocytes cultured with 20mM lactate, compared to the expression of cardiomyocytes without lactate (dashed line). The data shown corresponds to one representative isolation of mouse cardiomyocytes. Student’s T-test compared to control without lactate for each timepoint, *p < 0.05. **(B)** Relative expression of maturation genes in cardiomyocytes cultured with 20mM lactate, compared to the expression of cardiomyocytes without lactate (dashed line). The data shown corresponds to one representative isolation of mouse cardiomyocytes. Student’s T-test compared to control without lactate for each timepoint, *p < 0.05. **(C)** BMP10 protein quantification from cell lysate at different timepoints. Student’s T-test compared to control without lactate of the same timepoint, *p < 0.05.

Apart from the upregulation of progenitor genes promoted by lactate, we also investigated the expression of some genes associated with cardiomyocyte maturation. Glutamate receptors are excitatory neurotransmitter receptors that regulate cation influx and electrical impulse. Thus, we evaluated the expression of the glutamate receptor *Grik1* as a marker for mature and electrically active cardiomyocytes (Figures 3B and S3B). We observed that the expression of this gene was significantly reduced in the presence of lactate, especially during the first period of incubation. In addition, we analyzed the expression of the diacylglycerol kinase *Dgkk,* a key enzyme for lipid signaling and metabolism, characteristic trait of mature cardiomyocytes [11]. Lactate also downregulated the expression of *Dgkk* after 2 days of culture (Figures 3B and S3B).

To further confirm the immature cardiomyocyte phenotype, we assessed the production of protein BMP10 and, as shown in Figure 3C, protein levels were significantly increased on cardiomyocytes incubated with L-lactate compared to the control. Moreover, we used a cell cycle array to evaluate the expression of several genes key to cell cycle regulation (Figure S3C).

Even though these genes are predominantly regulated at protein level, we found higher levels of cell division cycle genes (*Cdc7, Cdc25c* or *Cdc20*) and genes associated to cancer and dedifferentiation (*P63, Ddit3, Mdm2, Sfn, Brca2* or *Ppm1d*). On the other hand, we could appreciate downregulation of some tumor suppressor genes and genes involved in cell cycle arrest (*Chek2, Cdkn2a* or *Atr*), further supporting the role of lactate on the promotion of an immature, proliferative and dedifferentiated state.

### *Ex vivo* hearts culture

Despite isolation and *in vitro* culture of cardiomyocytes is a versatile technique to study cardiac cells, it is an artificial environment far from the native conditions of the heart. For this reason, we studied the effect of lactate on whole neonatal mouse hearts cultured *ex vivo* (Figure 4A), maintaining the three-dimensional morphology of the heart while allowing contraction and cross-talking between different regions and cardiac cell types. Some parts of the heart tissue still showed spontaneous beating activity within the *ex vivo* culture period (Figure 4B). Around 20% of hearts in culture media without lactate (control) exhibited some beating abilities during the first 2 days, while none of the hearts cultured with lactate showed contraction. As expected, the number of control *ex vivo* hearts presenting beating regions decreased over time, probably due to lack of sufficient oxygen at the inner parts of the organ and cell death. Mouse hearts supplemented with lactate, however, exhibited increasing contractile abilities after 3 days of culture, with almost 90% of the evaluated hearts exhibiting beating regions at day 4 and around 60% after 7 days (Figure 4B).

**Figure 4.**
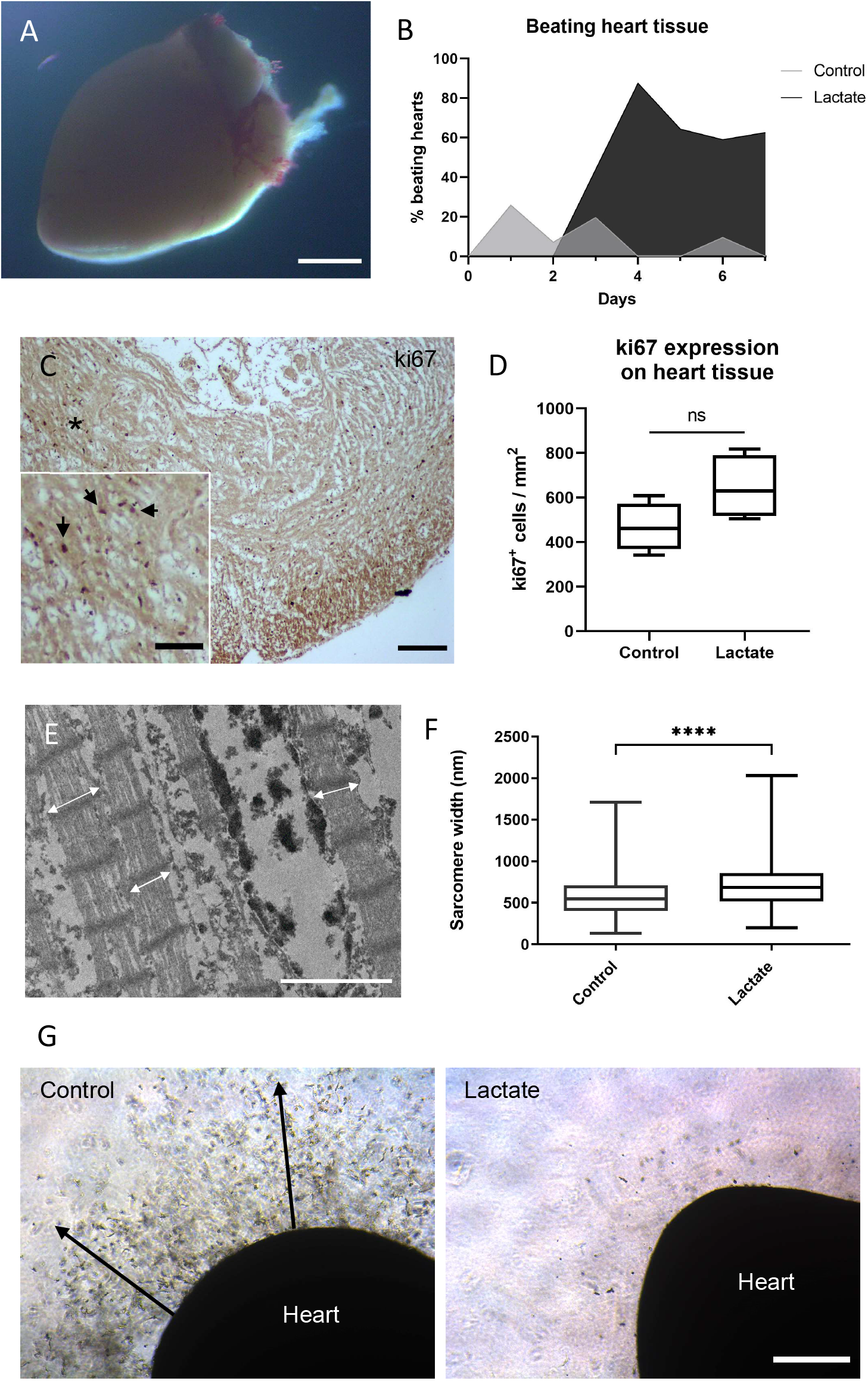
Ex vivo heart culture. **(A)** Stereomicroscope image of a cultured neonatal mouse heart embedded in Matrigel^®^ matrix. Scale bar: 1mm. **(B)** Fraction of ex vivo hearts showing spontaneous contraction (beating) activity at any region of the organ. 50 neonatal mouse hearts were evaluated. **(C)** Immunohistochemistry staining of ki67 on a cross-section of a neonatal mouse heart cultured ex vivo for 9 days. Black arrows indicate positive ki67 nuclei on a higher magnification image of the (*) region. Scale bar: 100μm and 50μm (magnification). **(D)** Ki67 positive cells quantification from immunohistochemistry performed on tissue sections from ex vivo hearts incubated with (Lactate) or without lactate (Control) for 9 days. Student’s T-test, ns: non-significant, p = 0.097, n = 4 hearts. **(E)** Transmission electron microscopy image of a cardiac tissue section from an ex vivo cultured heart after 4 days. White arrows indicate sarcomere width. Scale bar: 2μm. **(F)** Sarcomere width quantification from ex vivo hearts incubated with or without lactate for 4 days. Student’s T-test with Welch’s correction, ****p < 0.0001, over 800 sarcomeres measured from 4 hearts per condition. **(G)** Microscopy images of ex vivo hearts cultured with control (left) or lactate-supplemented media (right) for 9 days. Heart cells spread to the surrounding matrix on control hearts (black arrows). Scale bar 400μm.

Accordingly, cardiac cells from hearts incubated with lactate showed higher expression of the proliferation marker ki67 (Figure 4C and 4D) and a significant increase in sarcomere width, the functional unit of cardiac muscle contraction (Figure 4E and 4F). It was also interesting to observe that cardiac cells from control hearts appeared to leave the tissue structure after 9 days of culture and colonize the surrounding matrix, presumably escaping from the senescent *ex vivo* tissue. This event was not observed on lactate supplemented hearts (Figure 4G).

Altogether these results suggest that supplemented lactate can prevent the normal decline and senescence of the *ex vivo* heart tissue, which usually leads to tissue death and disorganization, and produce favorable conditions for cardiac cells to remain organized and keep their normal functions such as contraction.

### Effect of lactate on hiPSC-derived cardiomyocyte proliferation

Since our results demonstrated that lactate stimulates proliferation and dedifferentiation of neonatal mouse cardiomyocytes, we further explored the effect of lactate on a human model to verify whether it is conserved across these species. For this purpose, we used human induced pluripotent stem cell-derived cardiomyocytes (hiPSC-CM). First, we tested the viability of cells at different concentrations of lactate, from 0 to 20mM, using an MTS assay (Figure 5A). Lactate concentrations above 10mM demonstrated to reduce hiPSC-CM viability, showing up tolerance differences between mouse and human cells. Therefore, a lower range of concentrations up to 6mM was used to evaluate cardiomyocyte proliferation. Our results showed a concentration-dependent effect of lactate on the number of ki67^+^cTnT^+^ cells (Figure 5B) after only 24 hours of incubation (Figure 5C). Moreover, higher numbers of cardiomyocytes in karyokinesis were also observed (Figure S4A and S4B). After 4 days of culture, the positive correlation between lactate and ki67 expression was still observed (Figure 5D) (16.9 ± 2.7 % and 19.3 ± 8.4 % ki67^+^ cardiomyocytes for 2 and 6mM, compared to the control with 11.3 ± 4.1%; p = 0.037 and p = 0.006 respectively).

**Figure 5.**
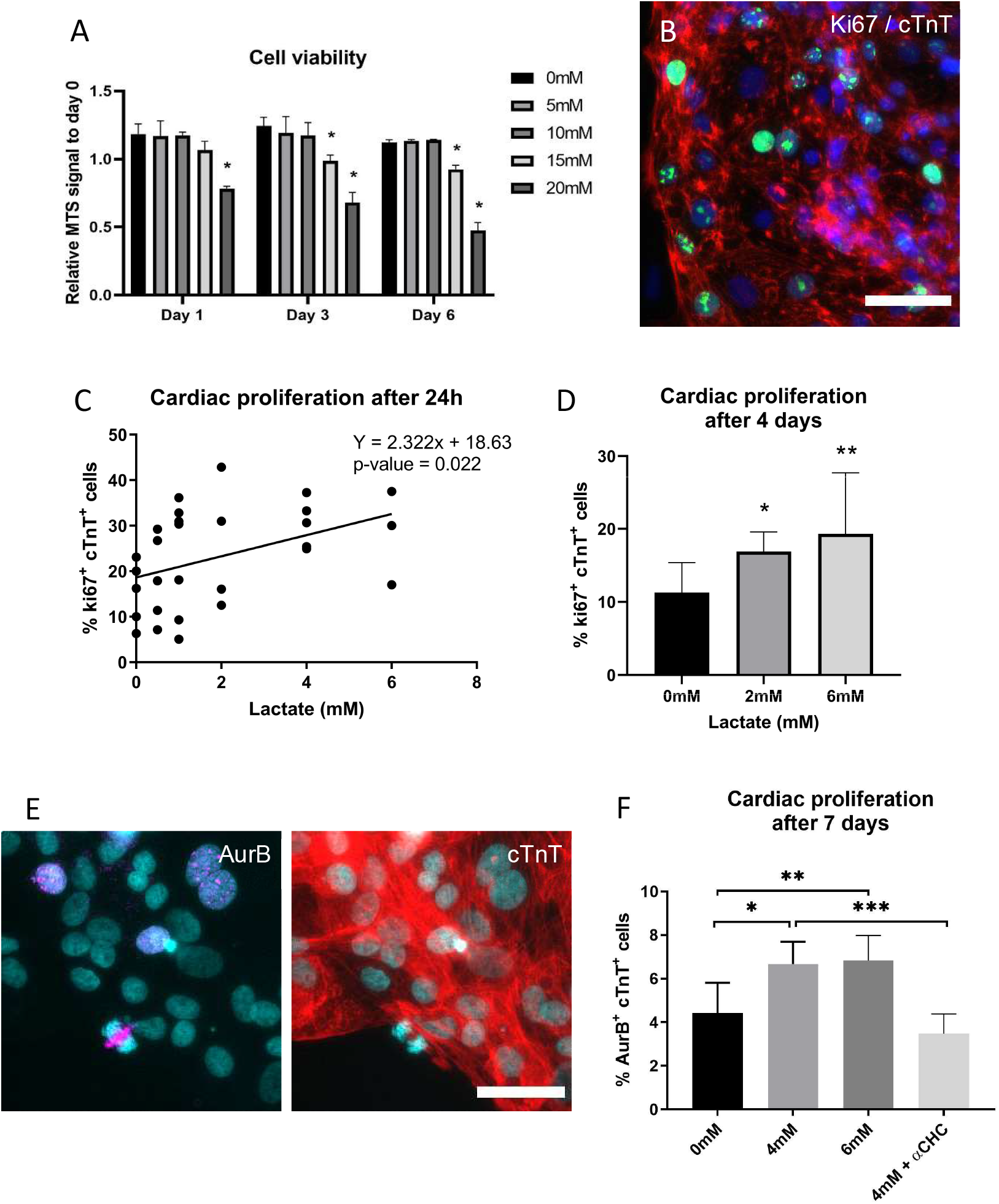
Effect of lactate on hiPSC-cardiomyocytes. **(A)** Cell viability in the presence of different concentrations of L-lactate (mM). MTS assay relative signal intensity is corrected to values from day 0 (1). Two-way ANOVA, *p < 0.01, n=3. **(B)** Immunofluorescent staining of ki67 on hiPSC-CM. Green: ki67, blue: nuclei, red: cTnT; scale bar: 100μm. **(C)** Ki67 quantification of positive cardiomyocytes from immunostaining after 24 hours of culture with different concentrations of L-lactate. Data are represented as % of ki67^+^cTnT^+^ within all cTnT^+^ cells. Simple linear regression, n ≥ 3. An average of 1500 cardiomyocytes were analyzed for each condition. **(D)** Ki67 quantification of positive cardiomyocytes from immunostaining after 4 days of culture with different concentrations of L-lactate. Data are represented as % of ki67^+^cTnT^+^ within all cTnT^+^ cells. One-way ANOVA, *p < 0.05, **p < 0.01, at least 8 random images were analyzed per condition. **(E)** Aurora-B immunostaining after 7 days of incubation. Red: cardiac Troponin T (cTnT), magenta: Aurora-B, cyan: DAPI; scale bar: 40μm. **(F)** Quantification of Aurora-B-positive cardiomyocytes after 7 days of incubation with 0, 4 or 6mM concentration of L-lactate. 1mM of α-cyano-4-hydroxycinnamic acid (αCHC) was used as inhibitor of MCT1. Data are represented as % of AurB+cTnT+ within all cTnT+ cells. One-way ANOVA, *p < 0.05, **p < 0.01, ***p < 0.001; at least 6 images and 4400 cardiomyocytes were analyzed per each condition.

To further validate the stimulation of cardiomyocyte proliferation and to discard possible polyploidy and polynucleation events, we studied the expression of Aurora-B kinase (Figure 5E). Consistently, the number of AurB^+^cTnT^+^ cells seemed to be positively correlated with L-lactate concentration after 2 days (Figure S4C), but this effect was significantly enhanced after 7 days of incubation with 4 and 6mM of lactate (Figure 5F). With the aim to identify the potential mechanism of action of lactate, we blocked lactate uptake into the cell using 1mM of α-Cyano-4-hydroxycinnamic acid (αCHC), a reversible inhibitor of the monocarboxylate transporter MCT1, the primary lactate transporter [27]. As shown in Figure 5F, the addition of αCHC successfully inhibited the increase in the number of AurB^+^ cardiomyocytes resulting from lactate supplementation. Therefore, we concluded that lactate uptake is an upstream requirement for triggering cardiomyocyte proliferation.

### Gene expression and transcriptomic analysis on hiPSC-CM

Gene expression analysis of hiPSC-CM exposed to 6mM of L-lactate revealed that *GATA4* was the main cardiomyogenic transcription factor affected by lactate supplementation (Figure 6A). Although *TBX5, NKX2.5* and *GATA4* are all critical regulators of cardiomyocyte lineage, they regulate different sets of genes. Consistently, *GATA4* is required for cardiomyocyte proliferation and the expression of cell cycle regulators [28], and it is a primary contributor to zebrafish heart regeneration [29]. Other cardiac markers, such as myosin heavy chain α (*MYH6*) and myosin light chain 2 *(MYL2),* predominantly expressed in cardiac atria and ventricle respectively, were not affected by lactate. Surprisingly, P63 was not affected on hiPSC-CM exposed to lactate, as well as the expression of *c-KIT*.

**Figure 6.**
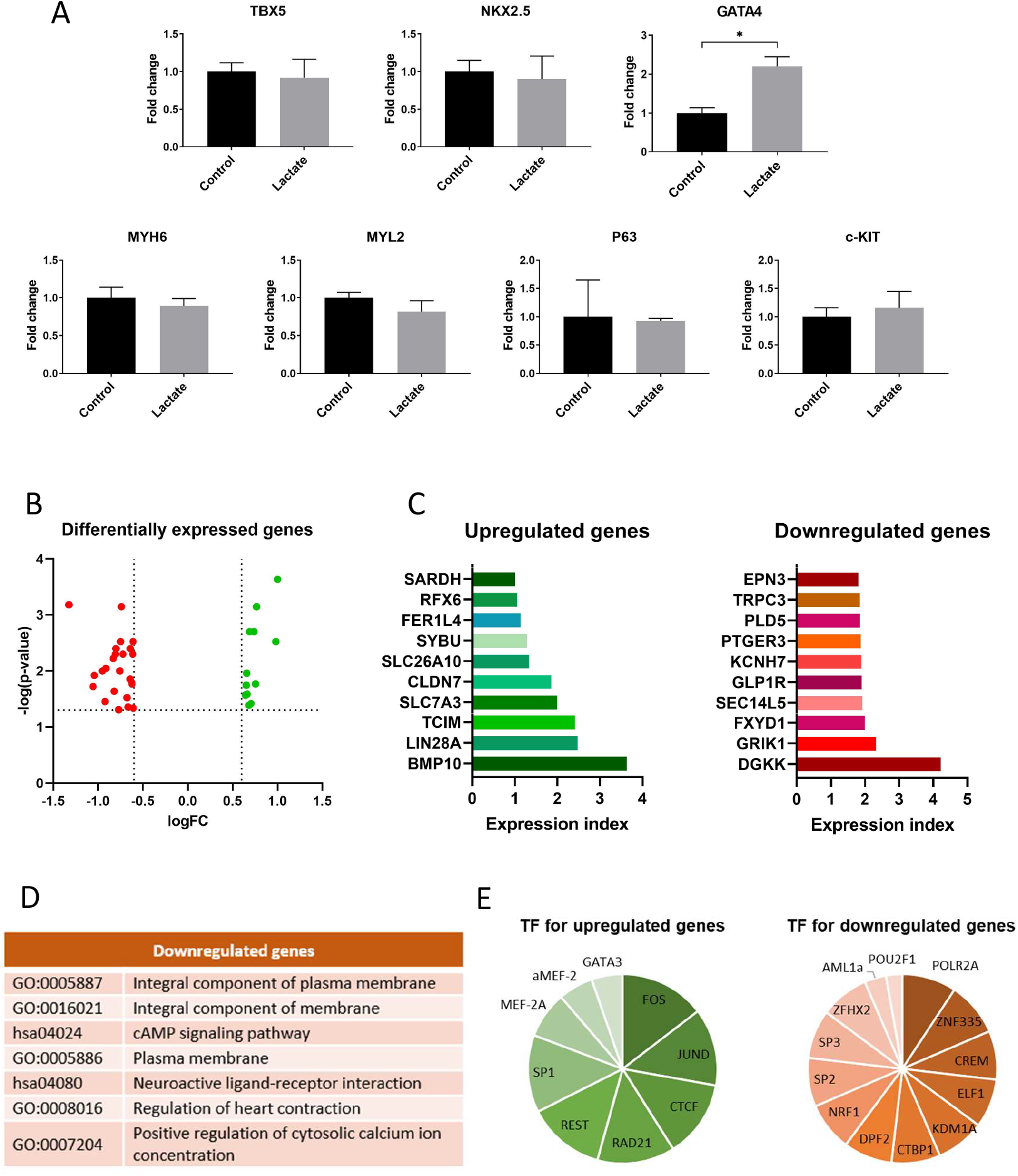
Gene expression and transcriptome analysis. **(A)** Relative expression of different cardiac genes with 0mM (Control) or 6mM lactate-supplemented media (Lactate) for 3 days. Student’s T-test, *p < 0.05. **(B)** Volcano plot showing differentially expressed (DE) genes from transcriptome sequencing. Dashed lines represent DE thresholds used (1.5 fold change cutoff and 0.05 p-value). Green: upregulated genes; red: downregulated genes; FC: fold change. **(C)** Top 10 up- and downregulated DE genes according to the expression index, the product of log(p-value) and logFC. **(D)** Gene ontologies and KEGG pathways associated to top downregulated genes (DAVID database, p < 0.05) **(E)** Top common transcription factors for upregulated and downregulated genes. All upregulated DE genes were used for the analysis, while only top downregulated genes were used. Data obtained from GeneCards and Qiagen databases.

To provide further insights into gene expression changes, hiPSC-CM were incubated with 6mM of L-lactate and analyzed via RNA sequencing. Transcriptomic analysis revealed a small number of differentially expressed (DE) genes (36 genes from 12388 genes with measured expression, using 1.5-fold change cutoff and 0.05 p-value), 12 upregulated and 24 downregulated genes (Figure 6B). This revealed a remarkably specific impact of lactate and the activation of a targeted molecular pathway. To simplify the analysis, an expression index was calculated for each gene as a result of the product of log (Fold change) and log (p-value). A list of the top 10 upregulated and downregulated genes by expression index is shown in Figure 6C. Genes associated to dedifferentiation and proliferation could be identified among the upregulated genes, specially *BMP10, LIN28A, TCIM* and *CLDN7*. Concerning the downregulated genes, we confirmed the repression of *GRIK1* and *DGKK* as the principal downregulated genes, but we could also identify some characteristic genes of mature cardiomyocytes, such as potassium (*KCNH7*) and calcium (*TRPC3*) ion channels. Gene ontologies (GO) and KEGG pathways associated to the most relevant downregulated genes (Figure 6D) included “cAMP signaling pathway”, “neuroactive ligand-receptor interaction”, “regulation of heart contraction” or “positive regulation of cytosolic calcium ion concentration” (DAVID database, p < 0.05). No common GO were found for the upregulated genes.

With the aim to shed some light into the regulatory mechanism by which lactate was exerting its effect on cardiomyocyte dedifferentiation, we evaluated the top common transcription factor binding sites found at the main regulatory elements of the DE genes (Figure 6E). Interestingly, most of these transcription factors were related to hypoxia pathways (Table S2 and S3). Some of them have already demonstrated important roles in cardiac hypoxia, such as *SP1, CREM, ELF1* or *MEF2A,* involved in cardioprotection or cardiac stem cell proliferation during hypoxia (Table S2 and S3). Overall, these data further corroborate a cardiomyocyte reprogramming produced by lactate towards dedifferentiation and proliferation, with hypoxia pathways suggested to be implicated on the underlying mechanism.

## DISCUSSION

The metabolic state of a cell is a key modulator of important biological processes, specially linked to pluripotency and stem cell differentiation [10,30]. The presence of fatty acids, the main substrate for adult cardiac metabolism, has already shown to induce maturation of cardiomyocytes [31]. In this context, this study demonstrates that lactate, a typical glycolytic metabolite, induces changes in gene expression leading to dedifferentiation and proliferation of mammalian cardiomyocytes. Apparently, this effect is restricted to the cardiomyocyte lineage, as neither cardiac fibroblasts (as shown in this work) nor undifferentiated stem cells [32] respond to the presence of lactate in a similar way. Tohyama *et al.* already showed distinct metabolic differences between cardiomyocytes and other proliferating cells, indicating the preferential use of lactate in cardiomyocyte metabolism [33]. Their study became the basis for a widely used approach to obtain enriched cardiomyocyte populations from human pluripotent stem cells. However, that approach is based on the elimination of stem cells in glucose-free conditions supplemented with lactate, thus obtaining an enrichment of the population of cardiomyocytes. In this work we have demonstrated that indeed, cardiomyocyte proliferation is stimulated by lactate, independently of glucose availability. These two events are complementary and may be occurring at the same time, as both processes require lactate uptake via MCT1. The ability of lactate to reprogram cardiomyocytes towards dedifferentiation and proliferation may have been masked until now by the more evident effect of glucose depletion on non-cardiomyocyte cells. This study has thus uncovered a crucial link between metabolism and dedifferentiation on mammalian cardiomyocytes.

We showed that lactate activates a specific genetic program inherent of cardiac progenitor cells. This reprogramming is driven by the expression of *LIN28, BMP10* and *TCIM* among others, along with the downregulation of *DGKK, GRIK1* and different ion channels. Importantly, Lin28 is a primal regulator of growth and metabolism in stem cells [8], while it also protects cardiomyocytes against apoptosis and regulates cardiac progenitor activation [34,35]. We have shown upregulation of BMP10, not only transcriptionally but also at a protein level. Bmp10 is a secreted protein that induces cardiomyocyte proliferation and prevents maturation of cardiac cells [25,36]. Thus, together with lactate, the production of Bmp10 may help to generate a pro-regenerative environment within the heart tissue, acting in a self-sustainable manner.

In addition, our results demonstrated that cardiac fibroblasts exposed to lactate reduce the production of inflammatory cytokines involved in apoptosis, hypertrophy or heart failure [37–39], while increase the expression of healing and regenerative cytokines [40,41]. Therefore, the synergistic response of cardiomyocytes and cardiac fibroblasts, the main cell types forming the heart, to the presence of lactate could potentially enhance heart regeneration by supporting new cardiomyocytes growth and survival. Our data are in concordance with results from *ex vivo* heart studies, where lactate inhibits epicardial cell spreading to the surrounding matrix (one of the major limitations of the *ex vivo* culture [42]) and promotes and maintains an immature and proliferative state on the three-dimensional native cardiac tissue, preventing tissue senescence and disorganization. These findings support the potential use of lactate in tissue engineering approaches, where it may mimic the early heart development [12] and play a crucial role on the maturation of cardiac tissue and the maintenance of an immature tissue state.

Neonatal mouse cell culture, however, has some limitations. The cardiomyocyte-enriched population of cells has a variable content of cardiac fibroblasts, with a rapid growth rate. This, together with an increase in cell death observed after 9 days of culture, hindered the effect of lactate at longer culture periods. This is especially evident when evaluating the expression of genes with a low basal expression level, such as *Dgkk,* and may be the cause for the decrease in the expression of *Tcim* after 7 days. Additionally, we found some differences on the effect of lactate between mouse and human cells. First, lactate concentration tolerance was higher in mouse neonatal cells than in hiPSC-CM. Secondly, *P63* was clearly overexpressed in mouse cells in the presence of lactate, while it was barely detected in human cells. This suggests that, even though *P63* has shown to play a critical role on cardiogenesis [24], their involvement in the lactate response may not be essential. Despite these differences, both cell types showed a similar response to lactate, suggesting the activation of a conserved pathway and making the development of a lactate-based therapy easily transferable for tissue regeneration therapies.

Nevertheless, the mechanisms of action of lactate are yet unclear. Some studies support the idea that lactate is directly affecting epigenetic modulation by affecting histone acetylation [43] or even by the direct lactylation of histones [44]. In this work, we suggest a common relationship between the observed changes in gene expression and important regulators of the hypoxic response, a hypothesis that is supported by the work of others [22,45]. Lee *et al.* proposed a molecular signaling pathway in which the accumulation of lactate due to hypoxic conditions activates Raf-ERK signaling pathways through NDRG3, subsequently promoting hypoxic cell growth [22]. Their results also suggest that the activation of this pathway can be induced at normoxic conditions by the addition of exogenous lactate, consistent with our findings. Additionally, it has been recently demonstrated that hypoxia is a key regulator of cardiomyocyte dedifferentiation and proliferation [46,47]. Taken together, lactate may be the key link of the cardiac response to hypoxia, as it is accumulated due to HIF1-mediated increase in glycolysis [22]. In the same way, lactate may explain the enhancement in adult cardiomyocyte proliferation caused by exercise [48] and observed after ischemic injury [49]. In these two events, an increase in cardiomyocyte lactate uptake has been reported [15,50].

The work presented here has been performed using neonatal and iPSC-derived cardiomyocytes, which differ from adult cardiomyocytes in their ability to proliferate and to exert a regenerative response [51,52]. But even though the adult heart is mainly comprised of post-mitotic cardiomyocytes, endogenous regenerative programs can be reactivated [4,53,54] and the presence of multipotent adult cardiac stem cells opens new opportunities for therapeutic myocardial repair [55]. The reactivation of Lin28a has already proven to enhance tissue repair in adults by reprogramming cellular bioenergetics [9], as well as to activate cardiac stem cells [35]. Furthermore, a population of hypoxic cardiomyocytes displaying proliferative regenerative potential have been shown to widely contribute to new cardiomyocyte formation in the adult heart [56]. Actually, it has been demonstrated that exposure to hypoxia induces a robust regenerative response in the adult heart, with decreased myocardial fibrosis and improvement of left ventricular systolic function [57]. These results are consistent with our findings regarding the effect of lactate on cardiac cells, further supporting the central role of this metabolite on hypoxia-mediated adult cardiac regenerative responses.

Therefore, the modulation of the metabolic microenvironment appears to be linked to key epigenetic and phenotype changes, even able to recapitulate other developmental stages such as the fetal growth. Either through the activation of cardiac progenitor cells or through proliferation of pre-existing cardiomyocytes, lactate supplementation has proven to induce cardiac tissue reprogramming to a fetal stage, where proliferation is crucial for organ growth and development. This opens new possibilities for the use of lactate as an easy and effective signal for the induction and maintenance of a regenerative program on mammalian cardiac cells. Furthermore, lactate is a common product from the breakdown of some biomaterials, such as polylactic acid and derivates. These biomaterials can be used as an effective lactate source for myocardial infarction or heart injuries, thus mimicking the production of lactate by placenta during fetal development.

Overall, this study has set the basis for the use of lactate and lactate-releasing biomaterials as metabolic modulators for new therapeutic strategies for cardiac regenerative medicine.

## ACKNOWLEDGEMENTS

We would like to thank Lourdes Sánchez-Cid and Sara Calatayud for animal care. Epifluorescence microscopy and image analysis was possible thanks to the Molecular Imaging Platform (MIP IBMB-PCB) and Dr. Elena Rebollo. We also thank CCiTUB for the help with sample preparation and TEM imaging, as well as for flow cytometry analysis; the Histopathology Facility (IRB) for heart tissue sample preparation; and MSU Genomics Core for sequencing services. We also wish to thank all members of the Engel and Aguirre Labs for valuable comments and advice.

## AUTHOR CONTRIBUTIONS

J.O. performed all experiments and data analysis. J.O., S.P. and E.E. designed and conceptualized all the experiments related to mouse cells. J.O. and A.A. designed and conceptualized the experiments concerning hiPSC. K.B. performed work on hiPSC culture and differentiation and sample preparation for RNA-sequencing. J.O wrote the paper. S.P. and E.E. supervised all the work.

## MATERIALS AND METHODS

### Isolation and culture of mouse cardiac cells

Cardiac primary cells were obtained from CD1 neonatal mouse following a modified protocol of Ehler, *et al* [58]. The procedure was approved by the Animal Experimentation Committee (CEA) from the Government of Catalonia. Hearts from 1-3-day-old mice were extracted and transferred on ice into a solution of phosphate-buffered saline (PBS) with 20mM 2,3-Butanedione monoxime (BDM, Sigma) where they were cleaned and minced into small pieces using curved scissors (approximately 0.5-1mm^3^ or smaller). Then, tissue fragments underwent a predigestion step by incubating in trypsin-EDTA solution 0.25% (Sigma) with 4μg/mL DNase I and 20mM BDM and subjected to 20-25 cycles of enzymatic digestion using collagenase II (Gibco) and Dispase II (Sigma) in L-15 medium (Sigma) with 20mM BDM. Pooled supernatants were collected through a 70μm nylon cell strainer (Corning) and centrifuged at 200G for 10 min The pellet was resuspended in DMEM containing 1g/L glucose (Gibco) supplemented with 19% M-199 medium (Sigma), 10% horse serum (Sigma), 5% fetal bovine serum (Sigma) and 1% penicillin and streptomycin (Gibco). Cell suspension was plated into a cell culture dish in order to separate the non-myocytic cell fraction of the heart, and after 1h of incubation the supernatant containing a purified population of cardiomyocytes was collected and centrifuged again. Cardiomyocyte cells were seeded on 0.5% gelatin-coated multiwell plates at 80-150 } 10^3^ cells/cm^2^ and cultured in DMEM containing 4.5g/L glucose (Gibco) supplemented with 17% M-199 medium (Sigma), 4% horse serum and 1% penicillin and streptomycin (Gibco). These cells consisted of an enriched population of beating cardiomyocytes, predominantly mononucleated. However, a minor population of non-myocytic cells were also obtained along with cardiomyocytes, irrelevant for experiments performed at short periods of time but significant when cells were cultured for longer periods, as non-myocytes presented a superior growth rate compared with cardiomyocytes.

Experiments without glucose were performed using DMEM without glucose (Gibco). The non-myocytic cell fraction was cultured in DMEM supplemented with 10% FBS (fetal bovine serum), 1% L-glutamine and 1% penicillin and streptomycin. L-(+)-lactic acid solution (Sigma) was supplemented to the culture media according to the experiment. Cell media was replaced every 2-3 days.

### iPSCs culture and cardiomyocyte differentiation

Human induced pluripotent stem cell lines iPSC-1 and iPSC-L2 were obtained from the Aguirre Lab. hiPSCs had been validated for pluripotency and lack of karyotypic abnormalities. hiPSCs were cultured in chemically defined growth media, Essential 8 Flex (Life Technologies), on growth factor-reduced Matrigel^®^ (Corning)-coated plates and routinely passed with Relesr (Stem Cell Technologies). For cardiomyocyte differentiation, 60-80% subconfluent hiPSCs were treated with Accutase (Sigma) and re-plated on a new Matrigel^®^-coated plate as a monolayer. Cells were differentiated to cardiomyocytes using CHIR99021 (Selleck Chemicals) and Wnt-C59 (Selleck Chemicals) in RPMI 1640 media (Life Technologies) supplemented with B27 (Life Technologies) with or without insulin following previously established protocols [59]. L-(+)-lactic acid solution (Sigma) was added to the cell culture media at differentiation day 15. For MCT1 inhibition, α-Cyano-4-hydroxycinnamic acid (Sigma) was added together with lactate. All cell lines were maintained in an incubator (37°C and 5% CO2) with media changes as necessary.

### *Ex vivo* heart culture

Neonatal mouse hearts were obtained and cultured following an adapted protocol [42]. Briefly, hearts from neonatal mouse were extracted and transferred to PBS with BDM. The excised hearts were then placed in a 24-well culture plate containing Matrigel^®^ solution (combined with cardiomyocyte culture medium 1:1). After gelation at 37°C, pre-warmed culture medium was added to each well containing a heart. Culture media was supplemented with 20mM of L-lactate and half volume of media was changed every 2-3 days. Beating movements of muscle tissue were evaluated daily under an inverted microscope. A cohort of 50 neonatal hearts was used.

### Transmission electron microscopy ultrastructural analysis

After 4 days of culture, mouse neonatal hearts were fixed in 2.5% glutaraldehyde and 2% paraformaldehyde in 0.1M phosphate buffer overnight at 4°C. They were rinsed in phosphate buffer and post-fixed with 1% osmium tetroxide (EMS) for 2 hours. Samples were then dehydrated in acetone series from 50 to 100% and infiltrated and embedded in Epon resin (EMS). Ultrathin sections of 60nm were obtained using a UC6 ultramicrotome (Leica Microsystems) and stained with 2% uranyl acetate and lead citrate. Sections were observed in a Tecnai^TM^ Spirit (FEI) transmission electron microscope equipped with a LaB6 cathode and images were acquired at 120kV with a CCD Megaviwe 1kx1k. Sarcomere width was measured using Fiji [60].

### Immunostainings

Cell samples were fixed in 4% PFA (EMS) and washed in cold PBS with glycine. They were permeabilized in 0.05% Triton^TM^ X-100 (Sigma) and blocked in 5% BSA or serum for 1 hour. Samples were incubated with primary antibodies against ki67 (Abcam ab16667 or ab15580), Aurora-B kinase (AurB or ARK-2) (Abcam ab2254), vimentin (Abcam ab24525), alpha smooth muscle actin (Invitrogen PA5-18292) or cTnT (Abcam ab8295) in 1% BSA overnight at 4°C. Secondary antibodies conjugated with AlexaFluor^®^ 488, 594 or 647 were used. Acti-stain^™^ Fluorescent Phalloidin (Cytoskeleton, Inc.) and DAPI (Sigma) were used as actin and nuclear counterstaining respectively. Images were taken using epifluorescence (Leica AF7000 and Olympus CKX53) and confocal microscopy (Zeiss LSM 800). Quantification and analysis were performed using Fiji.

Immunohistochemistry of neonatal mice hearts was performed after inclusion of hearts in paraffin. Samples were cut into sections of around 5μm using a microtome Leica RM 2155 and deparaffinized in xylene and ethanol, followed by antigen retrieval using citrate buffer at 95°C for 20 minutes. 4% goat serum (Abcam) in Tris-buffered saline containing 0.1% Tween^®^ 20 (TBST) was used as blocking agent and endogenous peroxidase was quenched with 3% hydrogen peroxide. Incubation with primary antibody against ki67 (Abcam) was performed overnight at 4°C, followed by incubation with biotinylated secondary antibody (Abcam ab64256), streptavidin peroxidase (Abcam ab64269) and DAB (Abcam ab64238). Images were acquired in a Nikon E600 and processed using Fiji.

### Real-time quantitative PCR

All reagents were obtained from Qiagen. RNA from mouse cells was extracted using the RNeasy^®^ Plus Mini Kit and cDNA synthesis was carried out using the RT^2^ First Strand Kit. The real-time PCR was performed with RT^2^ SYBR Green Mastermix using commercial RT^2^ qPCR Primer Assays and 18S ribosomal RNA as housekeeping gene. RNA from human cells was extracted using the RNeasy^®^ Mini Kit and reversed transcribed with the QuantiTect Reverse Transcription Kit. The QuantiTect SYBR Green PCR Kit was used for the RT-PCR, using commercial primers from IDT and Hprt1 as housekeeping gene.

### RNA-seq and transcriptomic analysis

RNA was extracted after lactate exposure at the indicated timepoints following RNeasy kit manufacturer’s instructions (Qiagen). Samples were processed using TruSeq RNA-seq sample prep kit from Illumina. Three independent samples per condition were loaded on an Illumina flowcell and clusters created by Illumina cBot. The clusters were sequenced in an Illumina HiSeq 4000 (MSU Genomics Core Facility). Read quality was assessed with FASTQC and reads were aligned using HISAT2 and the hg38 genome reference. EdgeR was used for differential gene expression analysis; values are reported as counts per million reads (cpm) or log2 FC. The data are available at the National Center for Biotechnology Information Gene Expression Omnibus repository under accession number GSE153938. Gene Ontology functional enrichment and clustering calculations were obtained from iPathwayGuide (Advaita Bioinformatics) and DAVID with a fold-enrichment of 1.5.

### Inflammation antibody array

Cardiac fibroblasts were incubated in 12-well plates in cell culture media containing 1% FBS and 0 or 20mM of L-(+)-lactic acid. After 24 hours, culture media was collected and cells were lysed in M-PER^®^ mammalian protein extraction reagent (Thermo Scientific). Cell media from 4 replicates was pulled as a single sample for analysis using a mouse inflammation antibody array (RayBio^®^, AAM-INF-1-8). Briefly, antibody array membranes were blocked and incubated with culture media samples overnight at 4°C. Then, membranes were incubated with biotinylated antibody for 2 hours at room temperature, followed by incubation with HRP-Streptavidin solution for 2 hours. Chemiluminescent detection was performed using an ImageQuant LAS4000 mini Biomolecular Imager (GE Healthcare Life Sciences). Intensity quantification was performed using ImageJ software and the obtained values were corrected by culture media background (without having been in contact with cells) and total protein content measured using the Pierce™ BCA Protein Assay Kit (Thermo Scientific). Values were further normalized by internal negative and positive controls from the antibody assay, as 0 and 100 expression values respectively. The experiment was repeated twice with independent non-myocytic populations coming from different mice isolations and results are shown as mean ± SD of these two independent experiments.

### Collagen quantification

Quantification of collagen was carried out by the colorimetric detection of hydroxyproline content. Briefly, cells were collected in M-PER Mammalian Protein Extraction Reagent (Thermo Scientific) and hydrolyzed in 6M HCl for 24 hours at 110°C. After hydrolysis, samples were dried and reconstituted in distilled water. They were then diluted in isopropanol and mixed with a Chloramine-T hydrate (Sigma) solution as oxidant agent and a 4-(Dimethylamino)benzaldehyde (Sigma) solution with perchloric acid as a color reagent. The absorbance was measured using an Infinite M200 PRO Multimode Microplate (Tecan).

### Wound scratch assay

Cardiac fibroblasts were cultured in 12-well plates for 7 days until confluence. A scratch of around 900μm was made using a sterile p200 micropipette tip. Cells were washed in PBS and incubated in DMEM media containing 5% FBS and supplemented with 0 or 20mM of L-(+)-lactic acid solution. Four images per well were taken along the wound using a Nikon TE200 microscope after the scratching and after 24 hours of incubation. Recovered area was calculated by measuring the scratched area of the same region before and after the incubation period.

Lactate toxicity was evaluated on primary mouse cardiomyocytes using calcein-AM (Sigma) and Propidium iodide (Fluka) for a live/death assay, and a Cytotoxicity Detection Kit^PLUS^ (Roche) for the detection of LDH. MTS Assay Kit (Abcam) was used to assess hiPSC cardiomyocytes viability.

### Cellular viability

Lactate toxicity was evaluated on primary mouse cardiomyocytes using calcein-AM (Sigma) and Propidium iodide (Fluka) for a live/death assay, and a Cytotoxicity Detection Kit^PLUS^ (Roche) for the detection of LDH. MTS Assay Kit (Abcam) was used to assess hiPSC cardiomyocytes viability.

### BMP10 ELISA

The presence of BMP10 on cell lysate was analyzed using a DuoSet^®^ ELISA Development System (Rs&D Systems, bio-techne).

### Statistical analysis

Data are presented as mean and error bars represent standard deviation (SD) from biological replicates (n). GraphPad Prism 8.3 was used as analysis and graphical software. Student’s T-test (two-tailed distribution) was generally used to compare two samples, and one-way ANOVA followed by post-hoc Tukey’s test was used for multiple samples, unless otherwise specified. A p-value < 0.05 was considered statistically significant.

