## Supplementary data and figures for "Lactate promotes cardiomyocyte dedifferentiation through metabolic reprogramming"

### SUPPLEMENTARY METHODS

**Cell cycle PCR array.** An RT2 Profiler PCR Array (Qiagen, PAMM-020ZC-2) was used for the detection of 84 genes related to cell cycle regulation following manufacturer's instructions. Only genes with a fold change above 1.2 or below 0.8 compared to control sample are shown.

**Cellular activity.** Mouse cardiomyocytes were cultured with different lactate concentrations and their activity was measured using Alamar Blue™ (Thermo Scientific). After 3 days of culture, cell media was removed and cells were incubated with 10% Alamar Blue™ in culture media for 2.5 hours, avoiding direct contact with light. Samples were collected in three replicates in a 96-well plate and the fluorescent signal (Ex: 528nm and Em: 590nm) was measured in a TECAN Infinite M200 Pro microplate reader. Sample values were normalized to the signal value of cells incubated without lactate (0mM).

**EdU incorporation.** EdU (5-ethynyl-2'-deoxyuridine) incorporation was used to evaluate cell proliferation, following EdU-Click 488 (Baseclick, BCK-EdU488-1) instructions. Briefly, cells were seeded at a cell density of around 40,000 cells/well in an µ-slide 8 well chamber (Ibidi) and incubated with 0, 5, 10 or 20mM of L-(+)-lactic acid solution (Sigma). After 24 hours, cell culture media from the chamber (in contact with cells) was mixed with the same volume of a 20µM solution of EdU solution in fresh medium (for a final concentration of 10µM of EdU) and added to cells. Cells were cultured in this new media (where lactate concentration was also diluted 1:2) for additional 24 hours. Cells were then fixed in 4% paraformaldehyde (EMS) for 15 minutes and washed twice in 3% BSA in PBS. Samples were permeabilized in 0.5% Triton X-100 in PBS for 20 minutes at room temperature. Afterwards, a reaction cocktail containing 6-FAM-Azide (prepared according to manufacturer's instructions) was added to samples, which were incubated for 30 minutes protected from light. Cells were finally washed in 3% BSA in PBS and counterstained with DAPI. Images were taken using a Leica DMIRBE microscope.

**Flow cytometry.** Cells were collected with trypsin-EDTA (Sigma) and fixed in 4% paraformaldehyde (EMS) for 10 minutes. Samples were then permeabilized with 0.05% TritonTM X-100 (Sigma) for 10 minutes and blocked with 3% BSA for 30 minutes. Conjugated primary antibodies anti-vimentin- AlexaFluor® 647 (Abcam ab194719) and anti-alpha-smooth muscle actin-FITC (Abcam ab188498) were used. A Beckman Coulter Gallios flow cytometer was used, where 13000 events were gated and analyzed from 2 biological replicates.

### SUPPLEMENTARY FIGURES AND LEGENDS

Figure S1. (**A**) Live/death assay of cardiomyocyte-enriched population of cells after 6 days of incubation with different concentrations of supplemented L-lactate. *Green*: alive cells; *red*: dead cells; scale bar 100µm. (**B**) Relative cellular activity of cardiomyocytes cultured with different concentrations of L-lactate (mM) for 3 days. One-way ANOVA, *p-value < 0.05. (**C**) Quantification of ki67-positive cardiomyocytes after 3 days of culture with (+) or without (-) glucose or lactate. Data are represented as % of $ki67^+cTnT^+$ within all $cTnT^+$ cells. One-way ANOVA, ***p < 0.001, ****p < 0.0001; 10 random images were analyzed for each condition. (**D**) Ki67-positive cells quantification from a cardiomyocyte population after 7 days of culture with 0mM (Control) or 20mM (Lactate) of lactate. In addition, lactate supplementation was removed from cell media after 3 days of culture (Lactate removal). Data are represented as % of ki67+ within all cells. One-way ANOVA, *p < 0.05; at least 6 random images were analyzed for each condition.

Figure S2. (**A**) EdU labelling of non-myocyte cells after 2 days of incubation with 5mM of L-lactate. *Blue*: nuclei, *red*: EdU; scale bar: 150µm. (**B**) Quantification of EdU-positive cells after 2 days of culture with different concentrations of L-lactate (mM). One-way ANOVA. 4 random images were analyzed for each condition. (**C**) Flow cytometry plots showing the (log) intensity of α-smooth muscle actin (α-sma) and vimentin of non-myocyte cells incubated with 0mM (Control) and 20mM (Lactate) of supplemented L-lactate for 4 days. The table shows the fraction of α-sma-positive cells among vimentin-positive cells calculated from 2 technical replicates of a single population of cells (Mean ± SD).

Figure S3. (**A-B**) Relative expression of progenitor (A) or maturation (B) genes in cardiomyocytes cultured with 20mM lactate, compared to the expression of cardiomyocytes without lactate at every timepoint (dashed line). The data summarizes different N isolations of mouse cardiomyocytes, being N ≤ 4. Student's T-test compared to control without lactate, *p < 0.05. (**C**) Relative expression of top up- and downregulated genes from a cell cycle array of cardiomyocytes incubated with 20mM of L-lactate for 6 days. Dashed line indicates the expression of genes from control cells without lactate.

Figure S4. (**A**) Detailed image of a cell nuclei in karyokinesis. *Grey*: nuclei; scale bar: 20µm. (**B**) Karyokinesis of cardiomyocytes ($cTnT^+$) quantified from DAPI staining after 24 hours. Approximately 3,100 cardiomyocytes were evaluated for each condition. (**C**) Aurora B quantification of positive cardiomyocytes after 2 days of incubation with different concentrations of lactate. Data are represented as % of $AurB^+cTnT^+$ within all $cTnT^+$ cells. One-way ANOVA. 7 random images were analyzed per each condition.

**Figure S1**

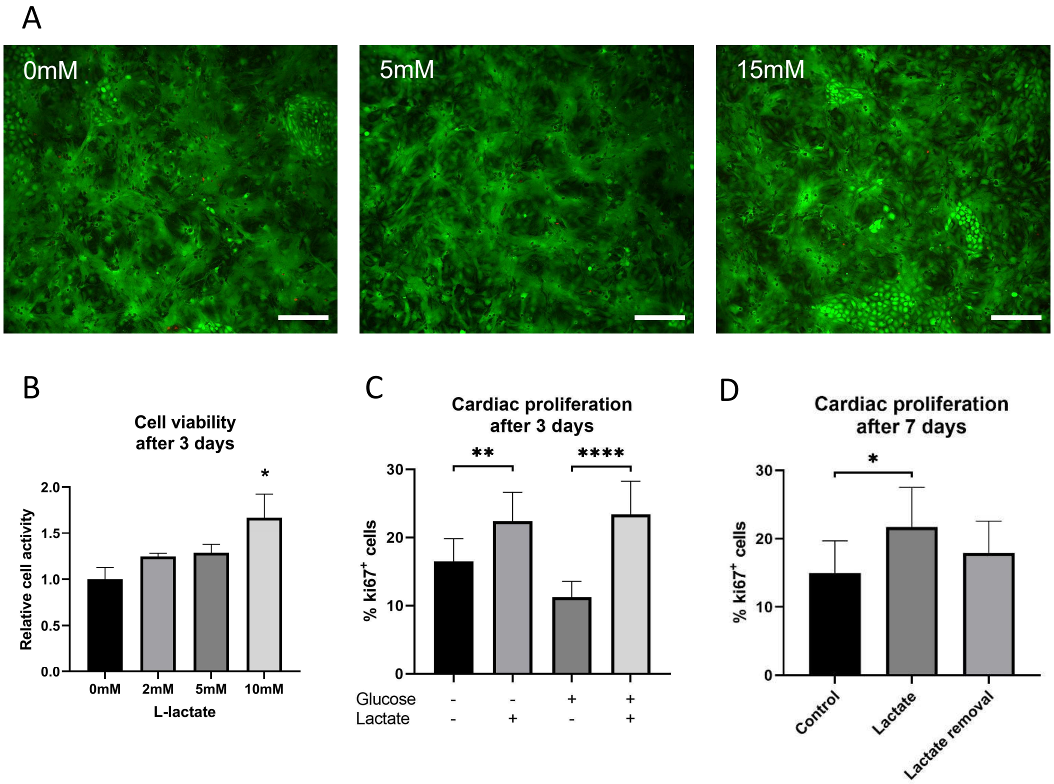

A
0mM
5mM
15mM
B
Cell viability after 3 days
Relative cell activity
2.0
1.5
1.0
0.5
0.0
*
0mM
2mM
5mM
10mM
L-lactate
C
Cardiac proliferation after 3 days
% ki67+ cells
30
20
10
0
**
****
Glucose - - + +
Lactate - + - +
D
Cardiac proliferation after 7 days
% ki67+ cells
30
20
10
0
*
Control
Lactate
Lactate removal

**Figure S2**

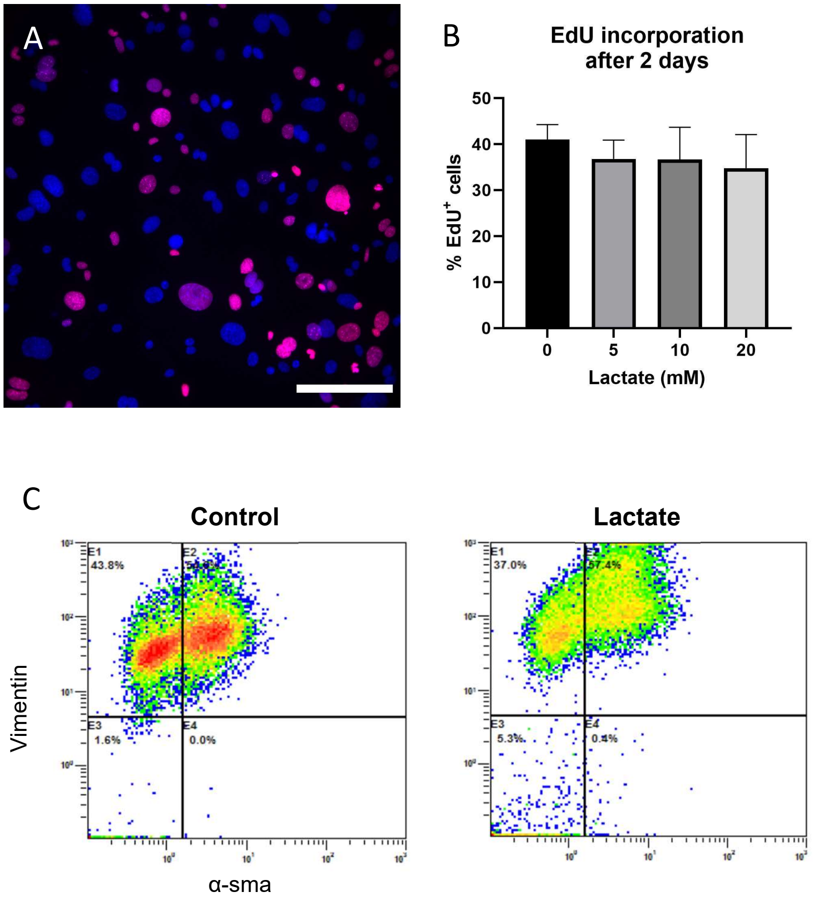


| | % α-sma⁺ vimentin⁺ cells |
|---|---|
| **Control** | 56,3 ± 1,17 |
| **Lactate** | 59,9 ± 1,34 |

**Figure S3**

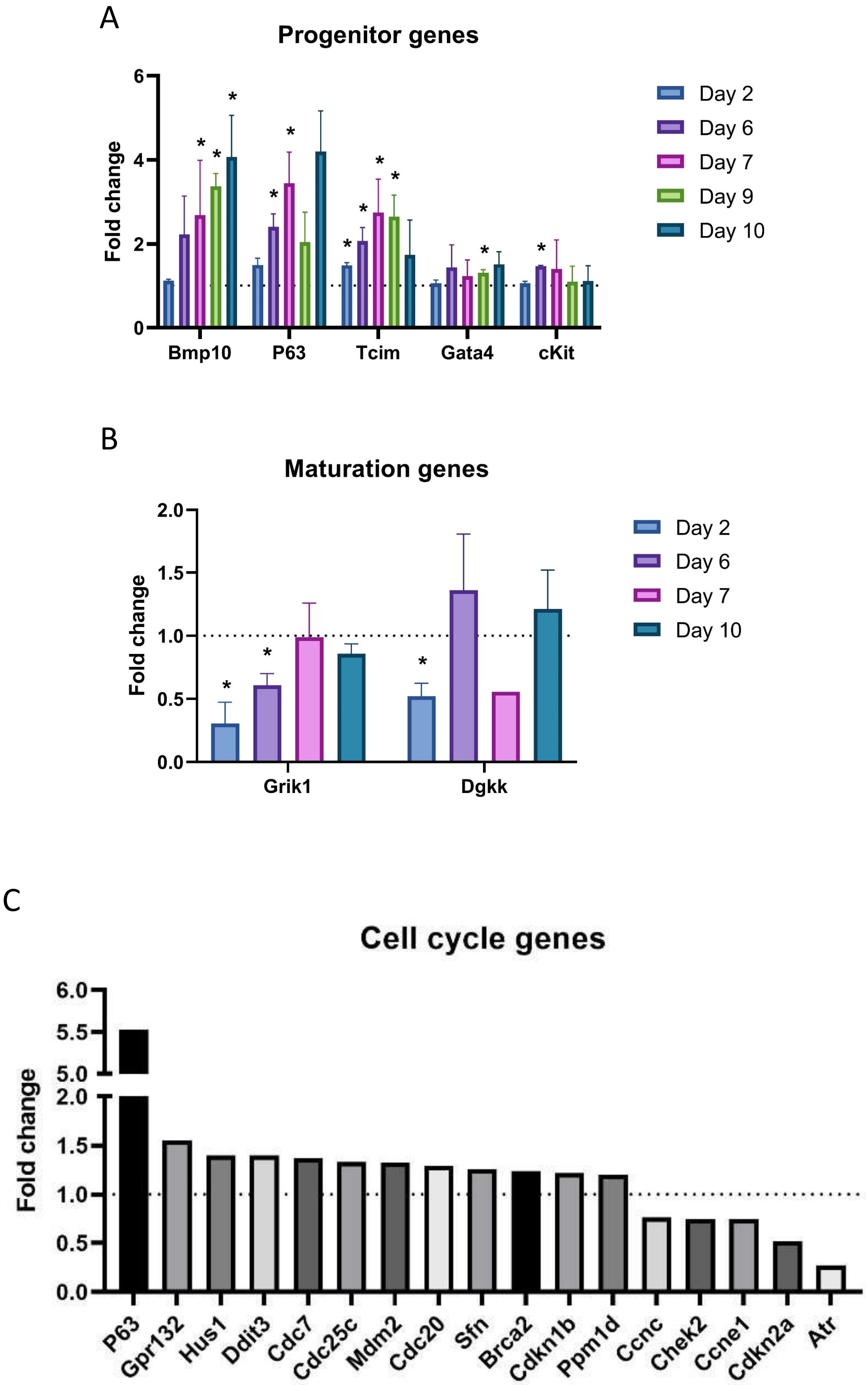
A
Progenitor genes
Fold change
Bmp10
P63
Tcim
Gata4
cKit
Day 2
Day 6
Day 7
Day 9
Day 10
B
Maturation genes
Fold change
Grik1
Dgkk
Day 2
Day 6
Day 7
Day 10
C
Cell cycle genes
Fold change
P63
Gpr132
Hus1
Ddit3
Cdc7
Cdc25c
Mdm2
Cdc20
Sfn
Brca2
Cdkn1b
Ppm1d
Ccnc
Chek2
Ccne1
Cdkn2a
Atr

**Figures S4**

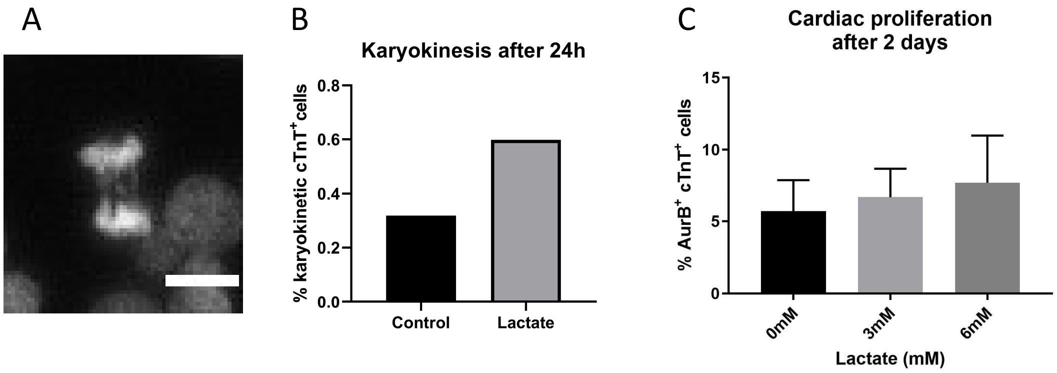

A
B
Karyokinesis after 24h
% karyokinetic cTnT+cells
0.8
0.6
0.4
0.2
0.0
Control
Lactate
C
Cardiac proliferation
after 2 days
% AurB+ cTnT+ cells
15
10
5
0
0mM
3mM
6mM
Lactate (mM)

### SUPPLEMENTARY TABLES AND LEGENDS

Table S1. **Inflammatory cytokines array**. Relative expression of inflammatory cytokines corrected by total protein amount and normalized to internal positive (100) and negative (0) controls. The mean and standard deviation (SD) was calculated from two independent cell isolations. Statistically significant differences (Student's T-test, p-value < 0.05) are indicated by an asterisk (*).

Table S2. **Transcription factors for upregulated genes**. Top common transcription factor binding sites for upregulated genes from RNA sequencing analysis. The up and downregulated genes presenting binding sites for the transcription factors on its main regulatory elements are shown, together with its relationship with hypoxia pathways. GeneCards and Qiagen databases were used.

Table S3. **Transcription factors for downregulated genes**. Top common transcription factor binding sites for top downregulated genes from RNA sequencing analysis. The up and downregulated genes presenting binding sites for the transcription factors on its main regulatory elements are shown, together with its relationship with hypoxia pathways. GeneCards and Qiagen databases were used.

Table S1

| Cytokine | Control mean | Control SD | Lactate mean | Lactate SD |
|---|---|---|---|---|
| **BLC** | 3.60 | 1.74 | 3.65 | 2.02 |
| **CD30** | 3.32 | 1.63 | 2.52 | 1.46 |
| **Eotaxin1** | 2.69 | 1.03 | 1.03 | 1.42 |
| **Eotaxin2** | 4.59 | 1.17 | 1.81 | 1.48 |
| **Fas** | 2.78* | 0.27 | -0.81* | 0.88 |
| **Fractalkine** | 5.40* | 0.81 | 0.26* | 0.53 |
| **GCSF** | 3.13 | 1.76 | -1.54 | 0.57 |
| **GM-CSF** | -1.18 | 1.33 | 0.29 | 0.16 |
| **IFN-g** | -0.24 | 0.00 | 0.71 | 0.55 |
| **IL1a** | 7.44 | 1.80 | 8.53 | 1.91 |
| **IL1b** | 1.16 | 0.85 | 2.03 | 1.17 |
| **IL2** | 2.04 | 0.95 | 2.92 | 1.40 |
| **IL3** | 2.57 | 1.70 | 3.23 | 1.63 |
| **IL4** | 5.53 | 2.50 | 5.84 | 2.24 |
| **IL6** | 13.07 | 5.50 | 15.11 | 5.85 |
| **IL9** | 5.64 | 1.52 | 4.22 | 1.39 |
| **IL10** | 2.32* | 0.22 | 0.08* | 0.28 |
| **IL12 p40/p70** | 3.40* | 0.81 | -0.09* | 0.03 |
| **IL12 p70** | 9.39 | 1.96 | 2.81 | 1.43 |
| **IL13** | 0.48* | 0.68 | 3.53* | 1.03 |
| **IL17A** | -0.14 | 0.14 | 1.21 | 0.84 |
| **I-TAC** | 2.04 | 0.95 | 2.42 | 1.11 |
| **KC** | 6.23 | 2.35 | 7.80 | 1.29 |
| **Leptin** | 2.04 | 0.95 | 3.01 | 0.29 |
| **LIX** | 14.95 | 5.33 | 18.85 | 6.19 |
| **XCL1** | 5.44 | 1.80 | 5.17 | 0.87 |
| **MCP1** | 28.94 | 11.52 | 33.76 | 4.98 |
| **MCSF** | 14.70 | 3.85 | 13.76 | 0.19 |
| **MIG** | 3.37 | 0.57 | 2.11 | 0.06 |
| **MIP1a** | 6.54 | 0.24 | 3.91 | 1.32 |
| **MIP1g** | 91.67 | 3.76 | 73.06 | 22.62 |
| **RANTES** | 1.33 | 0.89 | 3.88 | 1.53 |
| **SDF1a** | 1.09* | 0.46 | 4.67* | 1.41 |
| **I309** | 9.10 | 2.45 | 13.46 | 2.89 |
| **TECK** | 2.94 | 1.67 | 4.83 | 1.01 |
| **TIMP1** | 29.63 | 6.55 | 35.45 | 0.77 |
| **TIMP2** | 2.21 | 1.20 | 2.67 | 0.40 |
| **TNFa** | 1.44 | 0.96 | 1.78 | 0.62 |
| **TNF RI** | 14.30 | 1.58 | 11.00 | 3.89 |
| **TNF RII** | 12.89 | 2.42 | 11.58 | 3.06 |

### Table S2

| Transcription factors | Upregulated Genes | (Top) Downregulated genes | Relation to hypoxia |
|---|---|---|---|
| **FOS** | BMP10, LIN28A, TCIM, CLDN7, SLC26A10, SYBU, FER1L4, RFX6, SARDH, LINC00342, ADAMTS18 | GRIK1, SEC14L5, KCNH7, PLD5, EPN3 | [1,2] |
| **JUND** | BMP10, LIN28A, TCIM, CLDN7, SLC26A10, SYBU, FER1L4, RFX6, SARDH, ADAMTS18 | GRIK1, FXYD1, SEC14L5, PLD5, TRPC3, EPN3 | [1–4] |
| **CTCF** | LIN28A, TCIM, CLDN7, SLC26A10, SYBU, FER1L4, RFX6, SARDH, LINC00342, ADAMTS18 | GRIK1, FXYD1, SEC14L5, GLP1R, PLD5, TRPC3, EPN3 | [5] |
| **RAD21** | BMP10, LIN28A, TCIM, CLDN7, SLC26A10, SYBU, FER1L4, RFX6, SARDH, ADAMTS18 | GRIK1, FXYD1, SEC14L5, GLP1R, PLD5, TRPC3, EPN3 | [6] |
| **REST** | BMP10, LIN28A, TCIM, CLDN7, SLC26A10, SYBU, FER1L4, RFX6, SARDH, ADAMTS18 | GRIK1, FXYD1, SEC14L5, GLP1R, PLD5, TRPC3, EPN3 | [7] |
| **SP1** | LIN28A, TCIM, CLDN7, SLC26A10, SYBU, FER1L4, RFX6, SARDH, LINC00342, ADAMTS18 | DGKK, GRIK1, FXYD1, SEC14L5, GLP1R, PLD5, TRPC3, EPN3 | [2,8,9] |
| **CREB1** | LIN28A, TCIM, CLDN7, SLC26A10, SYBU, FER1L4 | GRIK1, FXYD1, SEC14L5, PLD5, EPN3 | [2,10–12] |
| **GATA3** | LIN28A, TCIM, SLC26A10, SYBU, | SEC14L5, GLP1R, KCNH7, EPN3 | [13] |
| **aMEF-2** | CLDN7, FER1L4, ADAMTS18 | GLP1R | |
| **MEF-2A** | CLDN7, FER1L4, RFX6, ADAMTS18 | GLP1R | [14–16] |

### Table S3

| Transcription factors | (Top) Downregulated Genes | Upregulated genes | Relation to hypoxia |
|---|---|---|---|
| **CREM** | DGKK, GRIK1, FXYD1, SEC14L5, GLP1R, PTGER3, PLD5, TRPC3, EPN3 | LIN28A, TCIM, CLDN7, SLC26A10, SYBU, FER1L4, SARDH, LINC00342, ADAMTS18 | [17] |
| **ELF1** | DGKK, GRIK1, FXYD1, SEC14L5, GLP1R, PTGER3, PLD5, TRPC3, EPN3 | BMP10, LIN28A, TCIM, CLDN7, SLC26A10, SYBU, FER1L4, SARDH, LINC00342 | [18] |
| **KDM1A** | DGKK, GRIK1, FXYD1, SEC14L5, GLP1R, PTGER3, PLD5, TRPC3, EPN3 | LIN28A, TCIM, CLDN7, SLC26A10, SYBU, FER1L4, RFX6, SARDH, ADAMTS18 | [19] |
| **CTBP1** | GRIK1, FXYD1, SEC14L5, GLP1R, KCNH7, PTGER3, PLD5, TRPC3, EPN3 | LIN28A, TCIM, CLDN7, SLC26A10, SYBU, FER1L4 | [20] |
| **DPF2** | GRIK1, FXYD1, SEC14L5, GLP1R, KCNH7, PTGER3, PLD5, TRPC3, EPN3 | BMP10, LIN28A, CLDN7, SLC26A10, SYBU, FER1L4, RFX6, LINC00342, | |
| **POLR2A** | DGKK, GRIK1, FXYD1, SEC14L5, GLP1R, KCNH7, PTGER3, PLD5, TRPC3, EPN3 | LIN28A, TCIM, SLC7A3, CLDN7, SLC26A10, SYBU, FER1L4, SARDH, LINC00342 | |
| **NRF1** | DGKK, GRIK1, FXYD1, SEC14L5, GLP1R, PTGER3, PLD5, TRPC3, EPN3 | LIN28A, CLDN7, SLC26A10, SYBU, FER1L4, SARDH | [21,22] |
| **SP2** | DGKK, GRIK1, FXYD1, SEC14L5, GLP1R, PTGER3, PLD5, TRPC3, EPN3 | LIN28A, CLDN7, SLC26A10, SYBU, SARDH | |
| **SP3** | DGKK, GRIK1, FXYD1, SEC14L5, GLP1R, PTGER3, PLD5, TRPC3, EPN3 | LIN28A, CLDN7, SLC26A10, SYBU, FER1L4, RFX6, SARDH | [2,23,24] |
| **ZFHX2** | DGKK, GRIK1, FXYD1, SEC14L5, GLP1R, PTGER3, PLD5, TRPC3, EPN3 | BMP10, LIN28A, CLDN7, SLC26A10, SYBU, FER1L4, SARDH, LINC00342 | |
| **ZNF335** | DGKK, GRIK1, FXYD1, SEC14L5, GLP1R, KCNH7, PTGER3, PLD5, TRPC3, EPN3 | LIN28A, CLDN7, SLC26A10, SYBU, FER1L4, RFX6 | |
| **AML1a** | GRIK1, FXYD1, GLP1R, TRPC3 | SLC7A3 | [25] |
| **POU2F1** | GRIK1, SEC14L5, PLD5 | SLC7A3 | [26] |
| **Pax-5** | GLP1R, EPN3 | FER1L4, ADAMTS18 | [27] |
